# Learning spatio-temporal V1 cells from diverse LGN inputs

**DOI:** 10.1101/2023.11.30.569354

**Authors:** Marko A. Ruslim, Anthony N. Burkitt, Yanbo Lian

## Abstract

Since Hubel and Wiesel’s discovery of simple cells and complex cells in cat’s primary visual cortex (V1), many experimental studies of V1 cells from animal recordings have shown the spatial and temporal structure of their response properties. Although numerous computational learning models can account for how spatial properties of V1 cells are learnt, how temporal properties emerge through learning is still not well understood. In this study, a learning model based on sparse coding is used to show that spatio-temporal V1 cells, such as biphasic and direction-selective cells, can emerge via synaptic plasticity when diverse spatio-temporal lateral geniculate nucleus (LGN) cells are used as upstream input to V1 cells. We demonstrate that V1 cells with spatial structures and temporal properties (such as a temporal biphasic response and direction selectivity) emerge from a learning process that promotes sparseness while encoding upstream LGN input with spatio-temporal properties. This model provides an explanation for the observed spatio-temporal properties of V1 cells from a learning perspective, enhancing our understanding of how neural circuits learn and process complex visual stimuli.

## 1 Introduction

The properties of simple cells and complex cells in the primary visual cortex (V1) were described by Hubel and Wiesel (Hubel and Wiesel, 1959, 1962, 1968). The properties of these cells can be characterized by their receptive fields (RFs). The visual RF of a neuron refers to the attributes of a visual stimuli that drive a neural response. For typical simple cells, their spatial RFs are spatially oriented, with alternating elongated ON and OFF subregions. Similarly, complex cells respond primarily to oriented edges and gratings, but unlike simple cells, they can display spatial phase invariance.

While the RF is often described solely in terms of spatial coordinates, the RF inherently involves both spatial and temporal dimensions. A RF is space-time separable if its three-dimensional structure can be written as the product of two independent functions: a two-dimensional spatial profile and a temporal profile. Otherwise, if a cell’s RF cannot be broken down into independent spatial and temporal components, its RF is space-time inseparable. Although numerous types of spatio-temporal inseparability are possible, simple cells with inseparable RFs show a highly characteristic profile in which the spatial phase of the RF shifts spatially over time. Experimentally, these cells exhibit direction-selectivity, namely that they are preferentially respond to motion of the visual stimuli in a particular direction (DeAngelis et al., 1995; De Valois et al., 2000). Non-directional V1 cells tend to have space-time separable RFs that are temporally monophasic or biphasic (DeAngelis et al., 1995; De Valois et al., 2000).

Simple cells in V1 receive upstream input from the lateral geniculate nucleus (LGN) and these LGN cells display diverse spatio-temporal properties. The spatial RFs of LGN neurons are typically unoriented with an approximately circular center-surround structure. Two different types of LGN cells display different temporal properties: Magnocellular LGN cells (M-cells) exhibit a biphasic temporal RF (i.e., their spatial RF will switch polarities over time), whereas parvocellular LGN cells (P-cells) are largely monophasic (i.e., their spatial RF will not change polarities over time). Due to these temporal properties, M-cells are sensitive to high temporal frequencies, tend to have transient responses, and thus are able to detect quick changes in visual stimuli, while P-cells are sensitive to low temporal frequencies and tend to have more sustained responses (Nassi and Callaway, 2009).

Previous experimental studies have suggested that both M-cells and P-cells contribute to the direction selectivity in V1 cells (De Valois et al., 2000; Saul et al., 2005). Various mechanisms have been proposed to explain the direction selectivity of V1 simple cells. One mechanism is that orientation-selective inhibitory neurons that have spatially offset receptive fields provide delayed input to excitatory neurons, which consequently leads to excitatory and inhibitory inputs that destructively interference in the non-preferred stimulus direction but not in the preferred stimulus direction (Freeman, 2021). Consequently, this asymmetry leads to direction selectivity. Another mechanism that could lead to direction selectivity is from the dynamic differences between ON and OFF LGN cells, namely differences in time delay and temporal kernel shape (Chariker et al., 2021). ON and OFF responses of LGN cells to sinusoidal gratings constructively or destructively interfere depending on stimulus direction. These models give possible theories of the mechanism of direction selectivity, but how the direction selectivity emerges from a learning process is still largely unknown.

Computational learning models have shown how the spatial properties of V1 cells can be learnt from static natural images using the principle of sparse coding (Olshausen and Field, 1996). There are biologically plausible models that can produce the diverse spatial properties of V1 cells (Zylberberg et al., 2011; Lian et al., 2019). However, these models fail to simultaneously produce the temporal properties that V1 cells possess. Recent work has shown how spatio-temporal properties of place cells in the hippocampus are able to be learnt from upstream spatio-temporal input using a model based on sparse coding (Lian and Burkitt, 2022). The goal of this study is to show that the spatio-temporal properties of V1 simple cells can be learnt using a model based on sparse coding. Namely, these diverse spatio-temporal properties of learnt V1 cells include the spatial properties of Gabor-like and blob-like receptive fields, temporal properties of spacetime separable RFs with monophasic or biphasic profiles, and the spatio-temporal properties of space-time inseparable RFs with direction selectivity.

## 2 Methods

### 2.1 Sparse coding

Our learning model of spatio-temporal V1 cells is based on the principle of sparse coding (Figure 1). The original sparse coding model was introduced by Olshausen and Field (1996), who proposed that the sensory input to each V1 simple cell can be represented by a small number of neurons. This model can be achieved by solving an optimization problem that reconstructs the input by a linear representation while also encouraging sparsity among model responses (Olshausen and Field, 1996, 1997).

**Figure 1.**
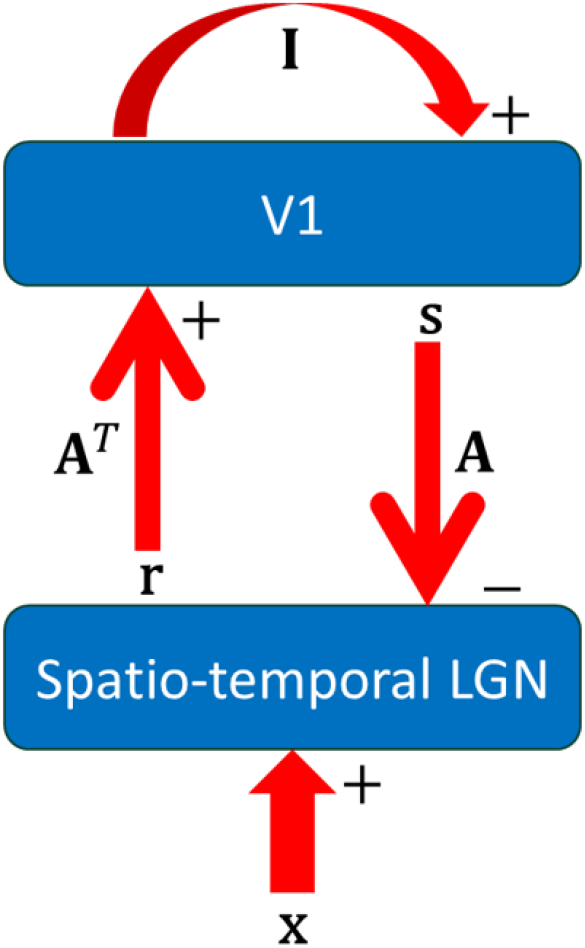
Model diagram. I is the identity matrix. **A**^*T*^ and **A** are plastic feedforward and feedback connections. **x** is the visual input.

Here, we implement the model based on the locally competitive algorithm that solves sparse coding efficiently (Rozell et al., 2008). The dynamics and learning rule of the model can be described by

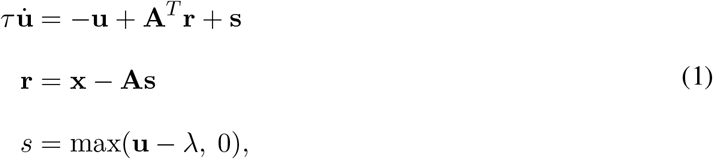

and

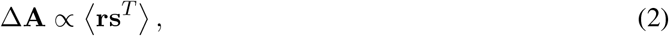

where *τ* is the time constant, **u** represents the membrane potentials of the model cells, **s** represents the non-negative firing response of the model cells, **r** = **x**–**As** is the reconstruction error, *λ* is a constant that determines sparsity, **A** are the synaptic weights and ⟨·⟩ denotes ensemble average.

### 2.2 Input

The input dataset used to train the model is a 2-minute video (1080×1920 pixels; frame rate 24 Hz) of natural scenes, which was taken in a forest walk (https://youtu.be/K-Vr2bSMU7o, with permission from the owner). For this study, this video is then trimmed to only keep the central 800×800 pixel region, similar to our previous study that learnt V1 complex cell properties (Lian et al., 2021).

#### 2.2.1 Spatial profile of LGN cells

Before the video is used for training, a pre-processing step is implemented to mimic the early visual processing done by LGN cells whose RFs have both spatial and temporal properties. The spatial properties of LGN RFs are characterised by a divisively normalized difference-of-Gaussian filters (Tadmor and Tolhurst, 2000; Ratliff et al., 2010):

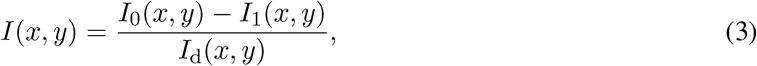

where *I*(*x, y*) is the spatially filtered pixel intensity at point (*x, y*), and *I*_0_, *I*_1_ and *I*_*d*_ are the outputs from unit-normalized Gaussian filters of center, surround and divisive normalization, respectively. The standard deviations of the center, surround and divisive normalization filters are chosen to be 1, 1.5 and 1.5 pixels, respectively. Eq. 3 is applied to each video frame to perform spatial pre-processing. Additionally, temporal filtering is applied across video frames to mimic the temporal properties of LGN cells, as described below.

#### 2.2.2 Temporal profiles of LGN cells

The temporal profile of the LGN RFs is either monophasic or biphasic. The LGN neurons with monophasic temporal profiles exhibit low-pass temporal frequency tuning, whereas the neurons with biphasic profiles have bandpass tuning (DeAngelis et al., 1995). We use axiomatically determined temporal models of LGN cells that satisfy time-causality and show qualitative agreement with biological receptive fields (Lindeberg, 2021). The monophasic temporal filter is created by convolution of truncated exponential functions *h*_exp_(*t*):

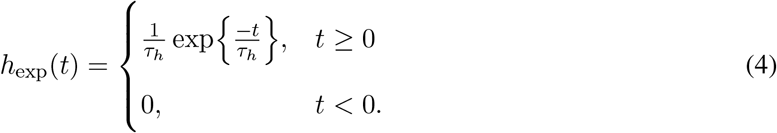

The convolution of *M* = 7 of these truncated exponential, scaled by a constant *k*_mono_ = 40, creates a temporally smooth monophasic profile, *H*_mono_(*t*):

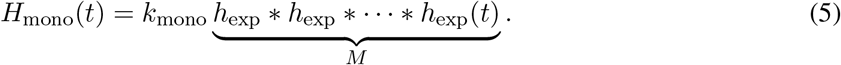

Temporal derivatives of this temporal model also satisfy the assumptions of a normative theory of visual receptive fields (Lindeberg, 2021). The first temporal partial derivative of this temporal profile multipled by a constant *k*_bi_ = 1.8 is used to define our biphasic temporal profile *H*_bi_(*t*):

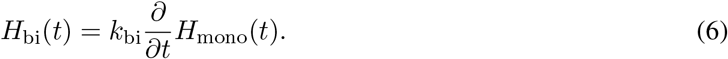

The monophasic and biphasic temporal profiles of the LGN used in this work are shown in Fig. 2. To apply these temporal profiles to the video stimulus, *τ*_*h*_ was chosen to be 15 ms, and the profiles are applied to the current and the 10 preceding video frames, evaluated at *t* = 0, 41.7, 83.3, …, 416.7 ms, since the frame rate of the video stimulus is 24 Hz. This yielded temporal profiles that qualitatively resemble those measured in LGN cells.

**Figure 2.**
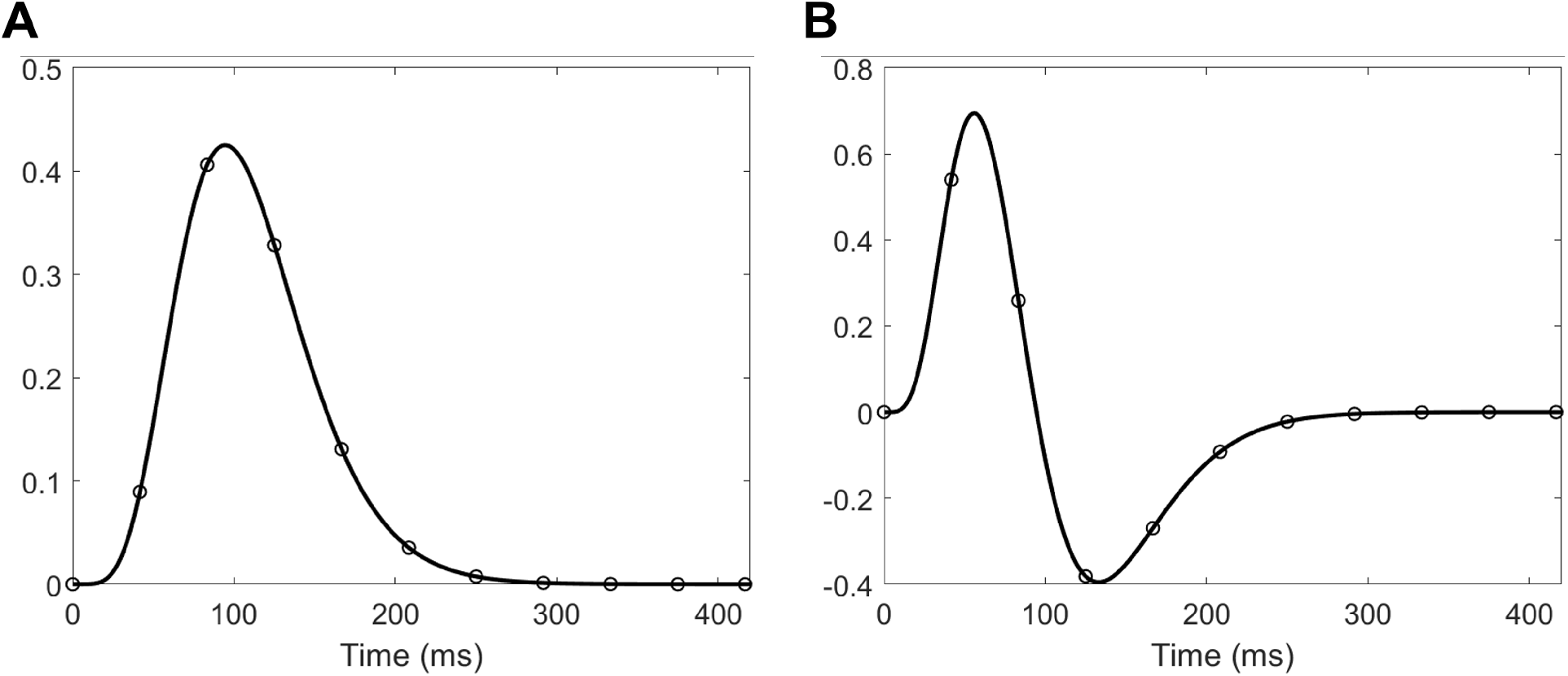
LGN temporal profiles. (A) Monophasic profile. (B) Biphasic profile. The open circle points represents the value of the temporal profile that is applied to the video frames.

In order to investigate how the temporal properties of LGN cells affects the temporal properties of learnt V1 cells, we define different scenarios where diverse LGN temporal profiles are used to temporally filter the input that has also been spatially filtered using Eq. 3.

**Scenario 1a** Modelled V1 cells only receive input from monophasic LGN cells. The LGN-V1 connection undergoes learning while the natural scenes video is presented. This scenario is designed to show that the spatial properties of V1 can be learnt, and also that their temporal properties can be inherited from LGN cells.

**Scenario 1b** Modelled V1 cells only receive input from biphasic LGN cells. Similar to Scenario 1a, the the LGN-V1 connection undergoes learning.

**Scenario 2** Modelled V1 cells receive input from both monophasic and biphasic LGN cells. The spatial and temporal properties of these LGN cells have been described previously. Similar to Scenario 1a & 1b, the LGN-V1 connection undergoes learning. This scenario is used to investigate whether diverse spatio-temporal properties of V1 cells can be learnt.

### 2.3 Training

Random video patches that have been spatially (Section 2.2.1) and temporally pre-processed (Section 2.2.2) are input to the model. Each frame of the video patch is of size 16 × 16 which results in 256 ON and 256 OFF LGN cells in the first layer. The number of model simple cells in the second layer is 256. Neural dynamics described by Eq. 1-1 were computed using the first-order Euler method. There are 50 integration steps with a time step of 1 ms.

The Hebbian learning rule in Eq. 2 is used to update the synaptic strengths. After each learning iteration, the columns of the weight matrix **A** are normalized to have an L2 norm of 1. Elements of this weight matrix **A** are initialized randomly from a Gaussian distribution and subsequently normalized before learning starts. For each learning iteration, 100 random 16 × 16 pixel regions of a video frame were presented.

The model was simulated in MATLAB (R2023b) and run on the high performance computing platform Spartan (Lafayette et al., 2016).

### 2.4 Recovering spatio-temporal receptive fields of learnt V1 cells

The method of spike-triggered average is used to estimate the receptive fields of V1 cells (Chichilnisky, 2001; Schwartz et al., 2006). We used white noise stimuli, **n**, of dimension 16 × 16 × *K* where *K* = 12 is the number of temporal frames to store. This white noise was spatially and temporally pre-processed (Sections 2.2.1 & 2.2.2). It was then presented to the model and the firing rate of a simple cell was recorded. Its spatio-temporal receptive field, **F**, which also has dimensions 16 × 16 × *K*, can be estimated through the weighted average,

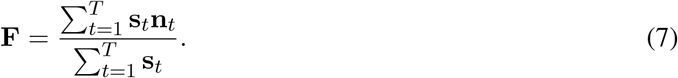

A way to visualise spatio-temporal receptive fields is by constructing an *x*-*t* plot (DeAngelis et al., 1995; De Valois et al., 2000). This plot summarises how the RF across one spatial dimension, *x* varies with time, *t*. The spatial dimension chosen for these plots is along the axis (noted as *x*-axis) perpendicular to the preferred orientation of the cell. Intensities of the RF across the spatial dimension perpendicular to *x*-axis were averaged to give the value at each *x* position.

### 2.5 Biphasic index

The biphasic index is a metric that quantifies the biphasic temporal characteristic of non-directional V1 cells (De Valois et al., 2000). For a non-directional cell, it is assumed that its RF is spatio-temporally separable, and so its temporal profile is extracted from its RF. The biphasic index is defined as the ratio, *I*_B_, of the amplitude of the second temporal peak, *P*_2_, to the amplitude of the first temporal peak, *P*_1_, of the temporal profile, where these two peaks have reversed polarity:

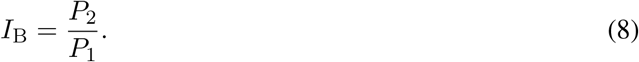

A low biphasic index close to zero suggests a temporally monophasic cell, and a high biphasic index indicates a very biphasic cell.

### 2.6 Direction tuning curve and directional index

The direction tuning curve is a response vs. direction plot that shows how the response of a neuron changes with different moving directions. To do this, we first search for the optimal spatial frequency of drifting sinusoidal grating that gives the maximal response of a model V1 cell. Then we present drifting sinusoidal grating with the optimal spatial frequency, at constant speed but of different moving directions (0 to 360^°^ with a step size of 1^°^).

To quantify a neuron’s degree of directional selectivity, we used the directional index (*I*_D_) that was used in experimental studies (De Valois et al., 2000):

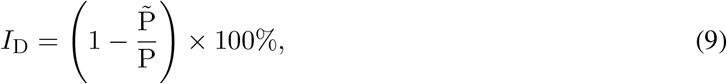

where P is the response in the preferred direction (i.e., the peak of the direction tuning curve), and 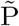 is the response in the non-preferred direction that is 180^°^ apart. As in De Valois et al. (2000), we defined direction-selective cells to be those with a directional index greater than 70, and non-directional cells to be those with a directional index less than 70.

## 3 Results

### 3.1 Learnt V1 cells have spatial RFs and monophasic temporal property if LGN cells incorporate monophasic temporal properties

After learning in Scenario 1a, the synaptic weights between the LGN input and model V1 cells shows good qualitative agreement with biological V1 receptive fields. We observe similar spatial structures such as oriented Gabor-like filters and non-oriented blobs. Similar to our previous study (Lian et al., 2019), we use the synaptic field to visualise the overall synaptic weights from ON and OFF LGN cells to V1, which is defined as

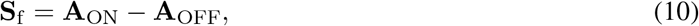

where **A**_ON_ and **A**_OFF_ represent the ON LGN to V1 connection and OFF LGN to V1 connection, respectively. As shown in Fig. 3A, the synaptic fields of V1 cells display receptive field shapes similar to biological V1 simple cells such as 2D Gabor filters and blobs.

**Figure 3.**
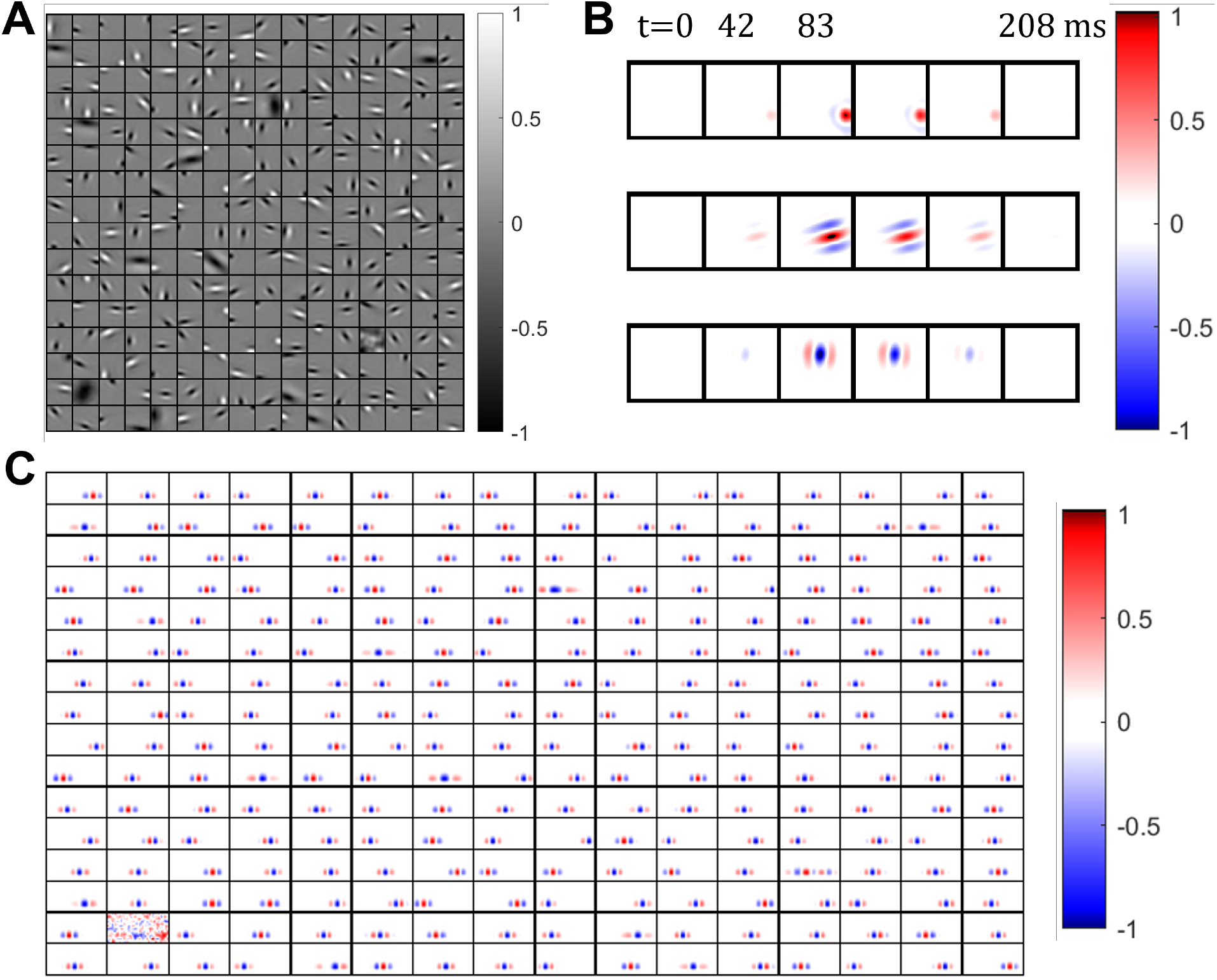
Receptive fields of V1 neurons only receiving monophasic LGN inputs. (A) Synaptic fields of all 256 model V1 cells visualized with spatial dimensions on the *x*- and *y*- axes. Values in each block are normalized to the range [-1 1]. (B) Dynamics of three example RFs at snapshots of video frame number or time, generated using STA. Values in each row are normalized to the range [-1 1]. (C) *x*-*t* plot of the spatio-temporal RF of all 256 model V1 cells, generated using STA, where the *x*-axis represents a spatial dimension and the *y*-axis represents time. Values in each block are normalized to the range [-1 1].

However, the synaptic fields only show a static representation of a cell’s receptive field while ignoring the temporal characteristics. Complete spatio-temporal receptive field profiles for representative model V1 cells are shown in Fig. 3B, where each row is a sequence of snapshots of a neuron’s spatial receptive field at different time steps. These neurons show a clear monophasic temporal profile, where the response builds up and then dies off over time. The peak appears at *t* = 83 ms, which is in agreement with the LGN monophasic temporal profile shown in Fig. 2A.

The spatio-temporal receptive fields of the whole population can be visualised by *x*-*t* plots (DeAngelis et al., 1995). The horizontal axis represents the spatial dimension perpendicular to that neuron’s preferred orientation, and the vertical axis represents time. All these *x*-*t* plots have a characteristic monophasic temporal profile. This is expected because the model V1 neurons only receive LGN input from monophasic cells. Additionally, the *x*-*t* plots show that these learnt simple cells have approximately space-time separable receptive fields.

### 3.2 Learnt V1 cells have spatial RFs and biphasic temporal properties if LGN cells incorporate biphasic temporal properties

For Scenario 1b, where the input LGN neurons have a biphasic temporal profile, the V1 neurons are also able to learn the diverse spatial features of receptive fields, namely oriented Gabor-like filters and unoriented blobs. The synaptic fields in Fig. 4A show the spatial representation of the model cells’ receptive field. A similar diversity of receptive fields can be seen compared to Scenario 1a. It should be noted that it took longer for these model cells to learn their receptive fields, since the input neurons only had a biphasic temporal profile. Naturally, this meant that the input neurons would have occasional and short-lasting activity, as is the case for M-cells in the LGN.

**Figure 4.**
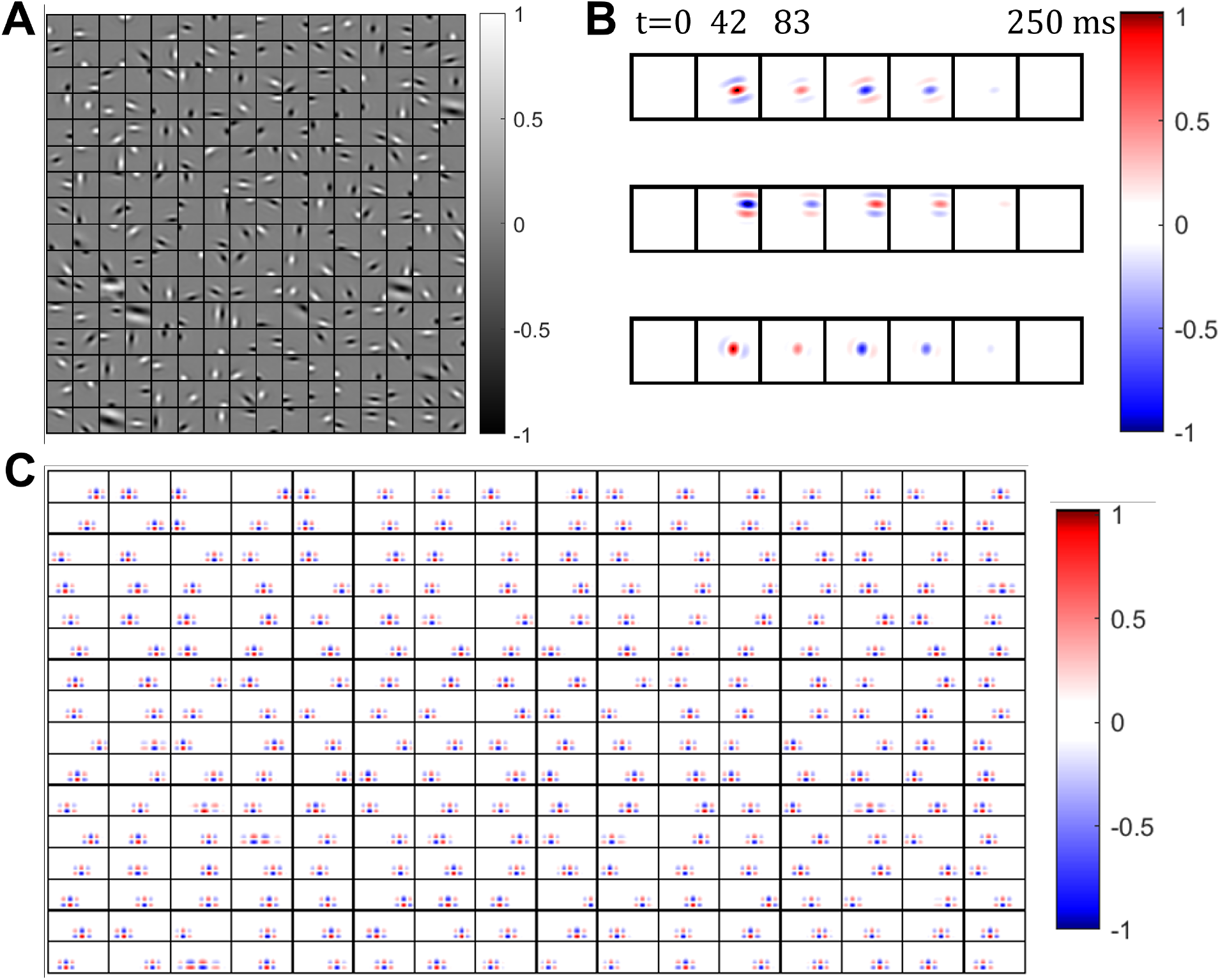
Receptive fields of V1 neurons only receiving biphasic LGN inputs. (A) Synaptic fields of all 256 model V1 cells visualized with spatial dimesions on the *x*- and *y*-axes. Values in each block are normalized to the range [-1 1]. (B) Dynamics of three example RFs at snapshots of video frame number or time, generated using STA. Values in each row are normalized to the range [-1 1]. (C) *x*-*t* plot of the spatio-temporal RF of all 256 model V1 cells, generated using STA, where the *x*-axis represents a spatial dimension and the *y*-axis represents time. Values in each block are normalized to the range [-1 1].

The time course of three example receptive fields are shown in Fig. 4B. Unlike the monophasic responses in Scenario 1a which rise then fall, the responses in this simulation show a biphasic profile that reverses in polarity. For example, the top cell initially has a central bright ON subregion surrounded by two dark OFF subregions above and below. After *t >* 83 ms, each subregion reverses polarity, so that the central region is now dark OFF. This follows from the LGN’s biphasic temporal kernel (Fig. 2B).

The spatio-temporal receptive field shown in Fig. 4C also highlights the biphasic property. Along the vertical axis, which represents time, the polarity reverses. All the model cells display a biphasic profile. Additionally, the biphasic temporal profile exhibited by the model neurons are essentially the same in terms of the amplitude of the temporal peaks, and the time course of the temporal profile, owing to the fact that only one type of temporal profile is used for the LGN inputs in this scenario.

### 3.3 Learnt V1 cells have spatial RFs and diverse temporal properties if LGN cells incorporate both monophasic and biphasic temporal properties

In Scenario 2, there are two LGN input populations. One population contains LGN neurons with a monophasic temporal profile, and the other population contains LGN neurons with a biphasic temporal profile. As a result, we produced two synaptic fields corresponding to these two LGN populations per model V1 cell, as shown in Fig. 5. Note that for some model neurons, most of the feedforward weights come solely from monophasic LGN inputs, while other model neurons primarily receive input from biphasic inputs and others receive input from both monophasic and biphasic LGN cells. However, for some of the model V1 neurons, the synaptic fields from monophasic LGN inputs and biphasic LGN inputs have their spatial phase slightly offset. Diverse spatial structures, in terms of receptive field location and orientation, can be seen from the synaptic fields, similar to the previous scenarios.

**Figure 5.**
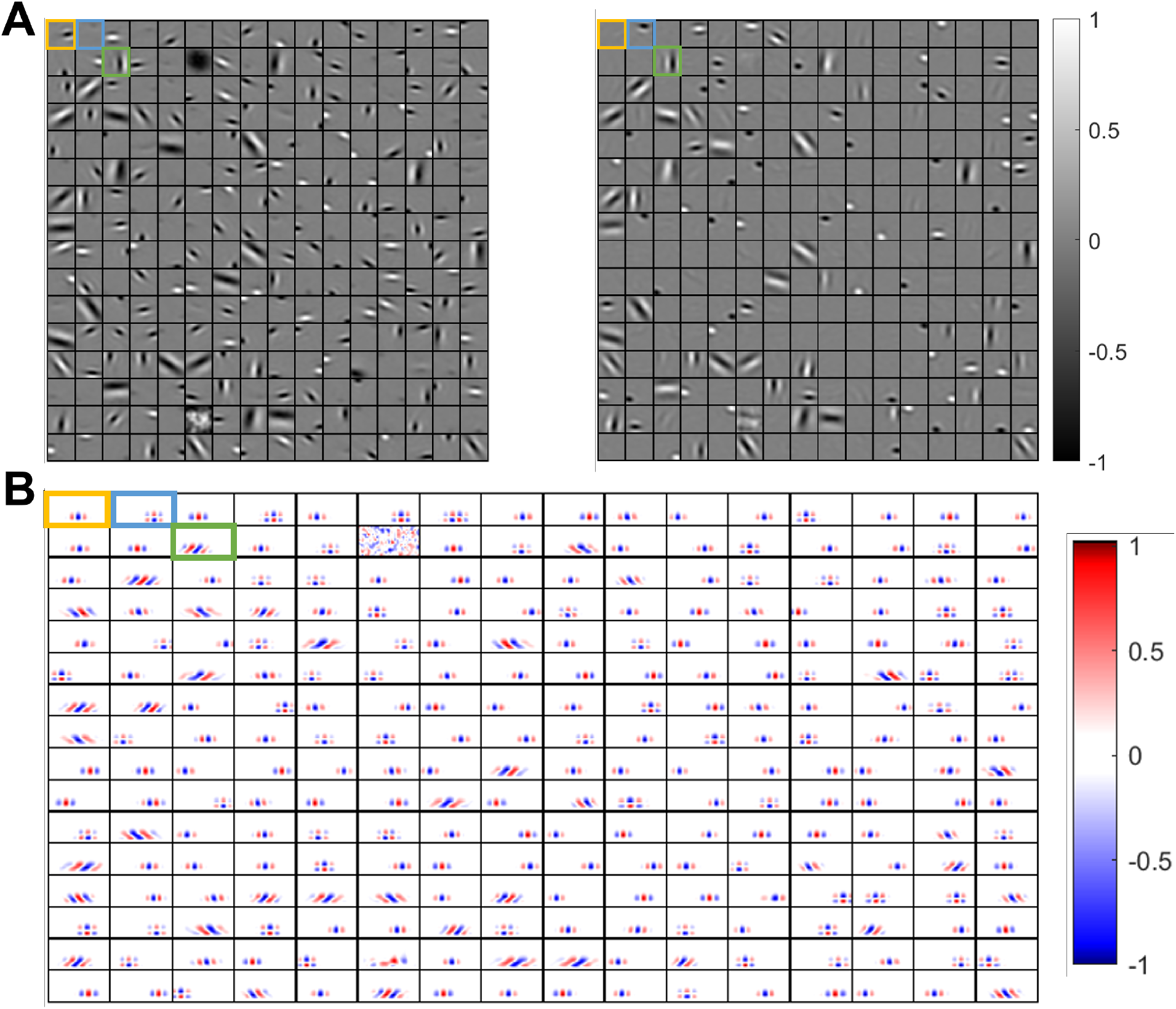
Receptive fields of V1 neurons receiving monophasic and biphasic LGN inputs. (A) Synaptic fields of all 256 model V1 cells from monophasic LGN inputs (left) and biphasic LGN inputs (right). Note that the *x*- and *y*-axes represent spatial dimensions. (B) *x*-*t* plot of the spatio-temporal RF of all 256 cells. Note that the *x*-axis represents a spatial dimension and the *y*-axis represents time. Yellow box highlights a model monophasic cell. Blue box highlights a model biphasic cell. Green box highlights a model direction-selective cell.

Unlike the previous scenarios, n the *x*-*t* plot shown in Fig. 5, a diverse range of temporal properties can be seen. There are some neurons that show a characteristic monophasic temporal profile and others that show a biphasic temporal filter. These neurons have separable spatial and temporal profiles. There are also neurons with receptive fields that are inseparable. In particular, their receptive field shifts in one direction over time. This can be seen by the slant in some of the *x*-*t* plots. This diversity in temporal properties is observed in experimental V1 neurons, where there are monophasic, biphasic or direction-selective cells.

Despite the LGN inputs not containing any motion-sensitive cells, V1 cells are able to learn direction-selective cells. An understanding for how this is possible can be seen in Fig. 6. The model cells that are direction-selective have inputs from both monophasic and biphasic LGN cells. However, these inputs are spatially offset by approximately 90^°^ in phase (Fig. 6A). Likewise, the temporal profiles of the monophasic and biphasic cells can be said to also have a 90^°^ phase difference. The combination of these spatial and temporal phase offsets of 90^°^ is termed spatio-temporal quadrature. The effect of the synaptic fields can be seen by convolving these synaptic fields with the LGN temporal profiles and visualising the result in an *x* − *t* plot (contour plot of Fig. 6B). The ON and OFF contributions from both monophasic and biphasic inputs create a striking slanted pattern. For illustrative purposes, we present estimates of the monophasic and biphasic contributions to the spatio-temporal receptive field (contour lines of Fig. 6B). Although the contour plots and the actual spatio-temporal receptive field do not perfectly align, the combined fields from the monophasic and biphasic contributions closely match the receptive field obtained by spike-triggered average.

**Figure 6.**
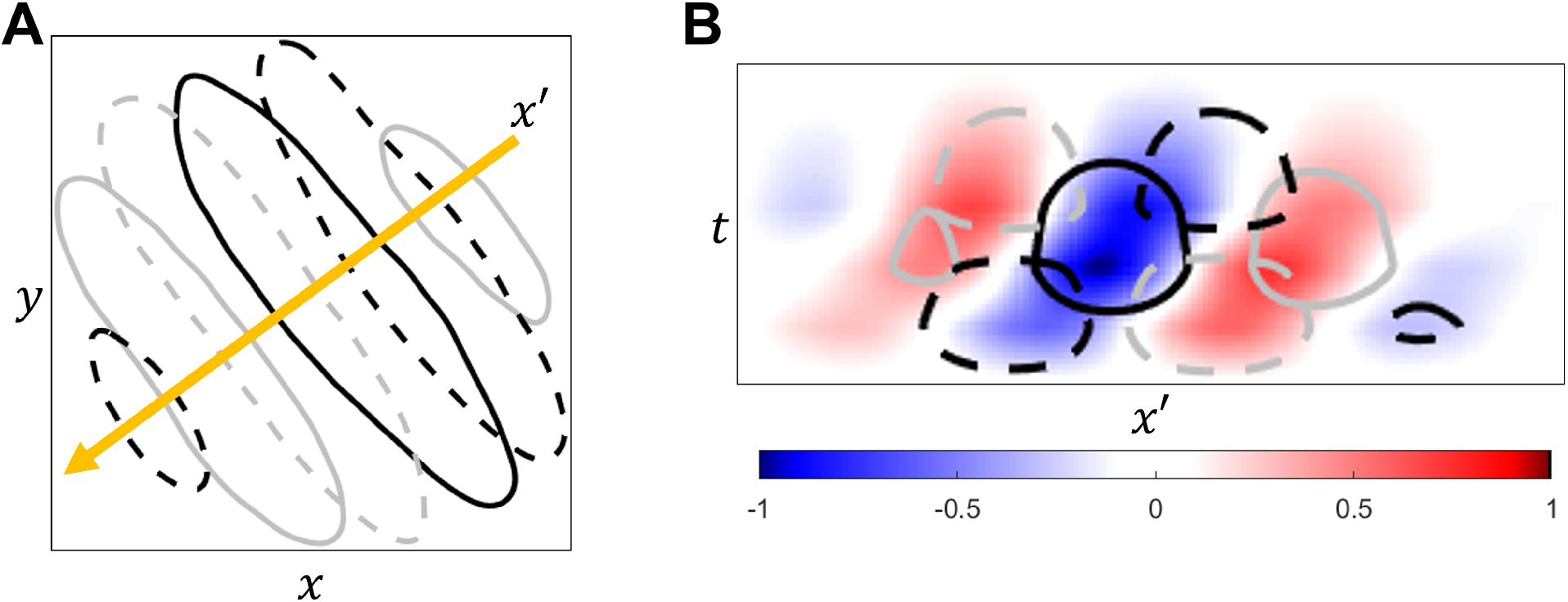
Direction selectivity arises from learnt monophasic and biphasic synaptic fields that are phase offset. (A) Monophasic (solid lines) and biphasic (dashed lines) synaptic fields for neuron 52 overlaid by a contour plot. Gray lines are ON regions and black lines are OFF regions. Yellow line is axis of the preferred direction of motion. (B) Spatio-temporal receptive field for the same model neuron with an estimate of the contribution from the monophasic and biphasic inputs plotted as contours. This estimate was calculated by convolving the synaptic fields with the LGN’s temporal filter.

Direction tuning curves were generated for each model cell by presenting sinusoidal gratings at each neuron’s preferred spatial frequency and in all directions. Four representative direction tuning curves are shown in Fig. 7A. The curve in blue shows an orientation-selective cell with a directional index of *I*_D_ = 0.1, where the neuron primarily responds in one orientation or two directions that are separated by 180^°^ with no preference for a particular direction. The curve in purple shows a very strong selectivity for direction, because the peak for the preferred direction is much greater than the peak for the non-preferred direction.

**Figure 7.**
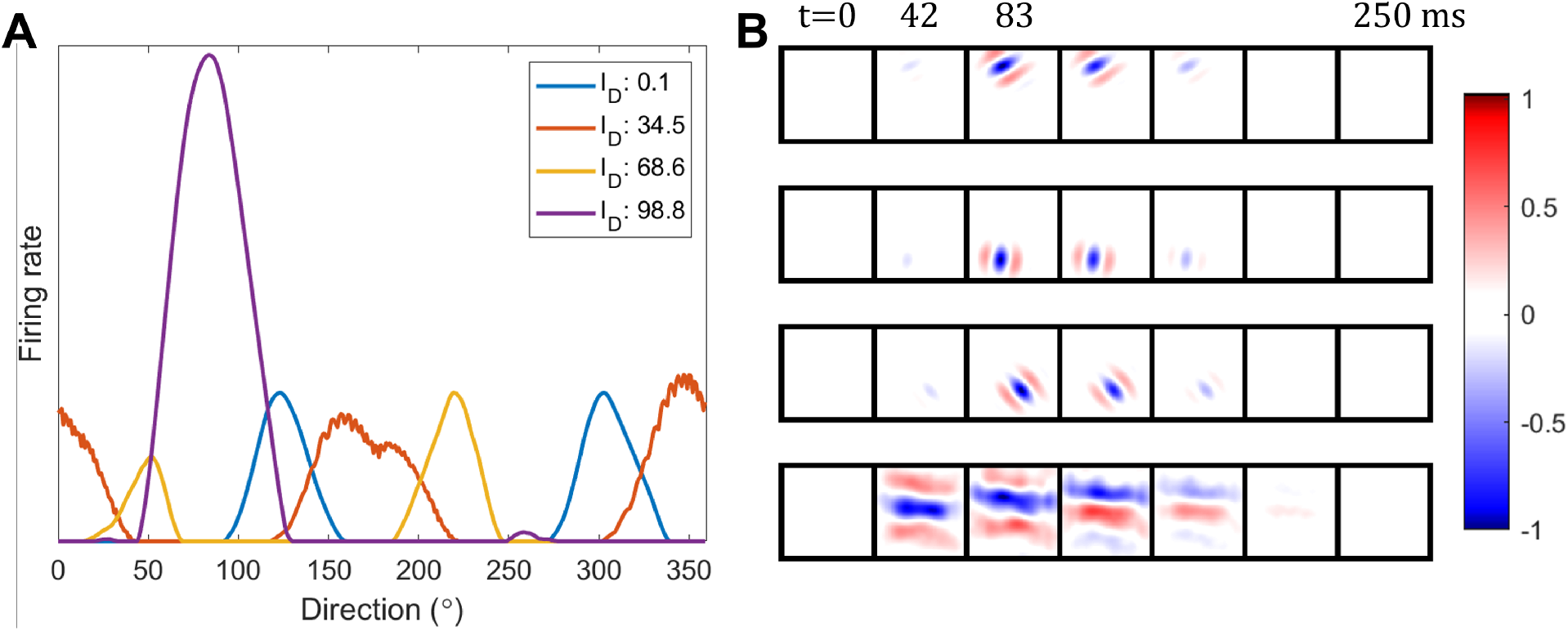
Direction tuning curves and receptive field dynamics. (A) Direction tuning curves of four selected model V1 neurons. (B) Dynamics of the four selected receptive field at snapshots of time, from least direction-selective (top) to most direction-selective (bottom).

Fig. 7B shows the time course of the receptive field of the neurons in Fig. 7. The top row corresponds to the orientation-selective cell with the blue direction tuning curve. This neuron prefers stimuli oriented at 135^°^ which is visible by its receptive field shape. The bottom cell corresponds to one direction-selective cell, with the purple direction tuning curve and a directional index of *I*_D_ = 98.8. This neuron prefers stimuli moving in one direction only and does not respond to stimuli moving in the opposite direction. From our simulations with both monophasic and biphasic input LGN cells, we show model V1 cells with diverse spatial and temporal properties. In particular, there are Gabor-like and blob-like spatial features, and there are neurons that are temporally monophasic, biphasic and direction-selective. From our simulations using these LGN properties, there are 207 direction-selective cells and 49 non-directional cells.

Fig. 8 shows the histograms of directional index of experimental V1 cells and learnt model V1 cells. As seen in Fig. 8A, there is a wide distribution of directional index in experimental data (De Valois et al., 2000).

**Figure 8.**
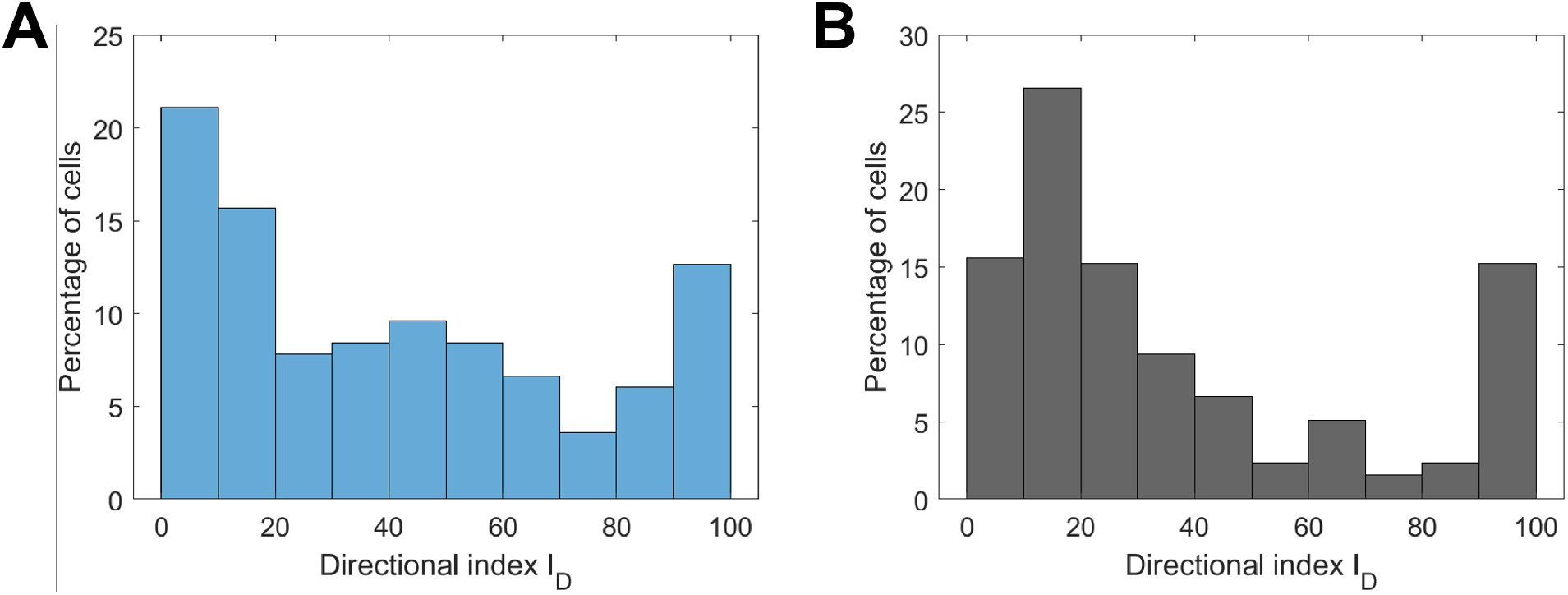
Distribution of directional index of V1 population. (A) Experimental data in macaque monkey (De Valois et al., 2000). (B) Model data after learning.

Their data is measured from 171 V1 cells in macaque monkey. The distribution of directional index of the model V1 cells is shown in Fig. 8B. The distributions of experimental and model are similar in that it is bimodal with the peaks occurring near zero, which represents non-directional cells and near 100, which represent direction-selective cells. However, other experimental data (such as Fig. 1 in Peterson et al. (2004)) suggests a different distribution for the same directional index. Their recordings of 310 simple cells are from adult cat visual cortex and their distribution exhibits an upward trend, with a significantly higher number of cells showing direction selectivity than those that do not. Nevertheless, the model V1 cells learn a diverse range of directional indices.

The biphasic index was calculated for each model non-directional cell (*n* = 207), as shown in Fig. 9. Similar to experimental data, we observe that the majority of non-directional cells are monophasic that have a very low biphasic index. In the data from De Valois et al. (2000), they found a diverse range of biphasic indices as well as a distinct small population of biphasic cells centred around *I*_B_ = 0.6 (blue histogram in Fig. 9A). In the model data, there is a similar peak at *I*_B_ = 0.6. However, this subpopulation is not as prominent in recordings by Saul et al. (2005) (red histogram in Fig. 9A) as well as in recordings in cat visual cortex (Peterson et al., 2004). Additionally, the distribution from model data appears to be trimodal with a lack of diversity seen in the distribution from experimental data. This could be due to the fact that the simulation in Scenario 2 only contains two types of LGN temporal profiles. LGN cells, however, display diverse temporal response properties, including lagged and non-lagged (DeAngelis et al., 1995), different time to peak responses and also different biphasic indices (De Valois et al., 2000). Discrepancies between experimental and model data will be further discussed in Discussion.

**Figure 9.**
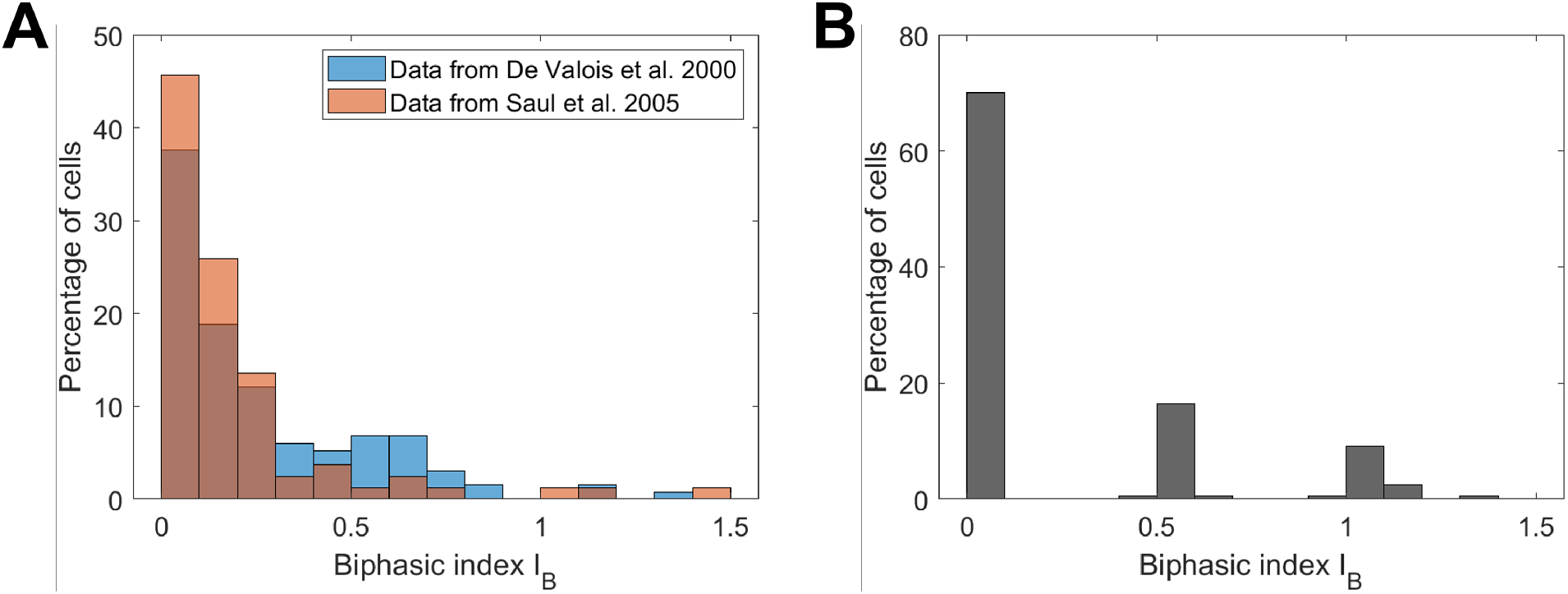
Distribution of biphasic index from the population of non-directional V1 cells. (A) Experimental data in macaque monkey from 134 non-directional cells reported by (De Valois et al., 2000) and 83 non-directional cells reported by (Saul et al., 2005). (B) Model data for 207 non-directional cells after learning.

## 4 Discussion

### 4.1 Summary

In this study, we use the sparse coding model to demonstrate that the temporal properties of V1 cells can be learnt simultaneously with spatial properties when upstream LGN cells have both spatial and temporal properties. Though V1 cells display diverse temporal properties such as biphasic response or direction selectivity, previous learning models of V1 cells primarily focus on the spatial properties. Our work shows that the principle of sparse coding can be used to learn spatio-temporal V1 cells from spatio-temporal LGN input, which suggests that both spatial and temporal properties of V1 cell can emerge as a result of cortical processing of upstream input. Combined with our previous work that demonstrates that spatio-temporal hippocampal place cells can be learnt from upstream spatio-temporal grid cells in the medial entorhinal cortex (Lian and Burkitt, 2022), it suggests that the processing of spatio-temporal upstream input may also account for the development of cell properties in other brain areas.

### 4.2 Discrepancies between model and experimental data

After learning, model V1 cells can display Gabor-like spatial receptive fields as well as diverse temporal properties such as monophasic response, biphasic response and directional selectivity. The two main measures used to compare model results with experimental results were directional index, *I*_D_, and biphasic index, *I*_B_. It should be noted that these measures have limitations, namely because they rely solely on two points of the response profile: the response at the preferred and non-preferred direction for the calculation of directional index, and the temporal peaks for the calculation of biphasic index (Saul et al., 2005; Mazurek et al., 2014). Relying on only two points of the response curve, rather than the entire profile, makes these metrics highly susceptible to noise. Other metrics, such as the circular variance for direction selectivity (Mazurek et al., 2014) and absolute phase for temporal properties (Saul et al., 2005) are more robust to noise, as they take into account the entire response profile. Additionally, the absolute phase can distinguish different types of transient responses that the biphasic index tends to group together (Saul et al., 2005). Nevertheless, the directional index and biphasic index measures were used in the study due to the availability of experimental data (De Valois et al., 2000; Saul et al., 2005; Peterson et al., 2004).

There are similarities in the general trend between the distributions of directional index and biphasic index of experimental data and model data. However, there are still some discrepancies (see Fig. 8 and 9). We suspect the differences may come from a more diverse LGN temporal properties in the real brain neural circuits. In this study, we incorporate both monophasic and biphasic temporal properties of LGN cells, but LGN temporal properties with the same type have exactly the same profile, which may lead to less diverse temporal input to V1 in our model than occurs in the brain. LGN neurons in the brain display a diverse range of temporal properties. These include non-lagged LGN cells, with a dominant first temporal phase, and lagged LGN cells, with a dominant second temporal phase (DeAngelis et al., 1995). As well as this, there are a range of values for the amplitudes and time-to-peak for the LGN inputs (De Valois et al., 2000; Peterson et al., 2004). Therefore, we infer that the model should learn more diverse temporal properties of V1 cells given heterogeneous LGN cells with a variety of temporal profiles.

### 4.3 Implementing the model in biologically plausible neural circuits

The sparse coding model used in this paper is implemented using locally competitive algorithm (Rozell et al., 2008), which is also used in other studies (Zhu and Rozell, 2013; Lian and Burkitt, 2021, 2022; Lian et al., 2023). Though some components of the model are not biologically plausible, the essence of this paper is to demonstrate that the principle of sparse coding can be used to account for the development of spatio-temporal properties in V1 cells.

A more comprehensive model may incorporate additional experimental findings. Experimental works propose that direction-selective simple cells are derived from the sum of two non-directional inputs which are also cortical simple cells, with a spatio-temporal phase difference of 90^°^, termed quadrature (De Valois et al., 2000; Peterson et al., 2004). The reason that these two inputs should come from V1 and not LGN is because the timing differences and temporal properties necessary to meet the conditions of quadrature are present in V1 (De Valois et al., 2000). However, the condition of spatio-temporal quadrature for producing direction selectivity can be relaxed (Peterson et al., 2004). Additionally, it is known that the monophasic and biphasic simple cells in V1 have their origins from the monophasic P-cells and biphasic cells M-cells in LGN respectively. Nevertheless, a more plausible circuit would be able to investigate the recurrence of non-directional simple cell connections to direction-selective simple cells.

Inhibition may play a crucial role in producing the direction selectivity and other temporal properties found in V1. Differences in the conductances of excitatory and inhibitory inputs in cat V1 were found when gratings were presented in the preferred and non-preferred direction (Priebe and Ferster, 2005). Inactivation of inhibitory neurons in ferret V1 reduced direction selectivity (Wilson et al., 2018). Nevertheless, the sparse coding model presented does not rule out contributions from inhibitory neurons, which could potentially be revealed by adapting the model to include a separate inhibitory subpopulation.

### 4.4 Conclusion

A biologically plausible model of V1 neural circuits would incorporate features such as spiking neurons that learn via spike-timing-dependent plasticity, a distinct inhibitory subpopulation, and recurrent connections. In our previous work, we developed such a model (Ruslim et al., 2024), and the next step is to integrate spatio-temporal LGN inputs as explored in this study. This enhanced model would allow us to test experimental findings more comprehensively, including the role of inhibitory neurons in shaping V1 response properties. Developing such a model that also aligns with the principle of sparse coding is an important and intriguing research question that forms part of our ongoing work.

## Acknowledgements

YL acknowledges funding from The University of Melbourne via 2023 Early Career Researcher Grant. ANB acknowledges funding from the Australian Government through the grant AUS-MURIB000001 associated with ONR MURI grant N00014-19-1-2571 and through the Australian Research Council’s Discovery Projects funding scheme (DP220101166).

